# Sociality changes Gene Essentiality

**DOI:** 10.64898/2026.06.23.734044

**Authors:** Emily Smith, Asia Humphrey, James Gurney

## Abstract

Transposon insertion sequencing (Tn-seq) has become a powerful tool for assigning gene fitness and essentiality in bacteria, and has recently been extended to bacteriophages. A core but rarely examined assumption of these screens is that fitness is measured in an asocial environment, where each mutant succeeds or fails on its own. Yet many genes act socially: their products can be shared among neighbors, allowing defective mutants to be complemented in trans. In phages multiple genotypes routinely coinfect the same cell. Here we show that social interactions distort gene essentiality. Using paired quorum-sensing microarray and Tn-seq data from *Pseudomonas aeruginosa*, we find that quorum-sensing–regulated genes are over-represented among genes scored as non-essential, confirming that social genes are under-reported as essential. We then build a stochastic, agent-based model of phage Tn-seq across an MOI gradient, assigning each gene an intrinsic fitness effect and a complementation fraction. Complementable (“social”) genes rise in frequency as MOI increases, masking their true fitness cost, whereas non-complementable (“private”) genes, do not. Partitioning genes by life-cycle stage and applying a two-round high-then-low-MOI design, further separates gene functions by life cycle stage. We argue that deliberate MOI manipulation turns a confound into a tool, enabling systematic classification of phage sociality.

## Introduction

Bacterio(phage) genomes are modular and densely packed with genes whose functions remain poorly understood ([1, 2]). Genome-wide mutagenesis approaches, such as transposon sequencing (Tn-Seq), have recently been developed for phages ([3, 4]). These pooled libraries assess the fitness of genes by tracking shifts in mutant frequencies before and after growth in a given condition ([3-5]). In bacteria, this framework has been used primarily to identify essential genes and map conditional dependencies across environments ([5-7]). However, some genes in these experiments are likely enriched due to social interactions. This issue is largely ignored in bacteria due to the scale of libraries in which ∼10^9 bacteria are all in competition with each other. This highlights an underlying assumption in the literature: gene essentiality is often determined under the assumption of an asocial environment. It is currently unclear if phage Tn-seq (or bacterial TN-seq for that matter) under reports gene essentiality due to social complementation.

In phages, the interpretation of pooled fitness assays is laid bare by the intracellular nature of viral replication. Because multiple phages can infect the same host cell, genes and their products may be shared in trans, allowing defective mutants to be complemented by coinfecting genotypes ([8-11]). Now instead of 10^^9^ genomes all competing and complementing each other with public goods as few as two might be in a single environment. To combat the issue of complementation between phages, researchers have used very low numbers of phages to limit the number of phages per bacteria to single phage infections ([3]). Higher phage infections per bacterial cell may result in Tn-Seq experiments that do not reflect asocial gene essentiality, but instead reflect the social environment experienced within infected cells. This means that Tn-seq, with careful consideration for MOI manipulation, might be an appropriate way to study the degree of phage sociality. Importantly, the likelihood of bacterial cells being infected by more than a singly phage is high [12, 13]; therefore, the social environment may actually be the more relevant ecological scenario upon which we should be making our assumptions when we study phage biology. Phage genes might have evolved under social conditions.

We argue that Tn-Seq in phages can be used to identify not only essential genes, but potentially also socially interacting genes, whose fitness effects depend on coinfection. Multiplicity of infection (MOI), here defined as the ratio of phage to bacteria in the initial experimental conditions and not the ratio of actual phage infecting any cell, can act as a tunable parameter that alters the scale of competition/cooperation. At MOIs below 1.67 (1.67 phage per bacterium), most infected cells are infected with phage propagules originating from a single phage (assuming a Poisson distribution), creating an essentially “asocial” environment where any competition or cooperation between phages only occurs between closely related replicates of the initially infecting phage (Fig 1). A mutant phage cannot be rescued by another coinfecting genome if none are present. At MOIs higher than 1.67, most bacteria are infected by at least two phages. Systems such as superinfection exclusion systems can increase the MOI required to have more than two phage per cell ([14, 15]), or alternatively, collective infection can reduce the MOI required to successfully infect a cell ([16]). For the sake of simplicity, we will ignore these possibilities in this work and return to it in the discussion. Coinfection promotes both complementation and exploitation, generating dynamics analogous to cooperation and cheating in bacterial social evolution ([16-20]). By measuring mutant frequency trajectories across MOI gradients, we could partition phage genes according to their degree of sociality.

**Figure 1.**
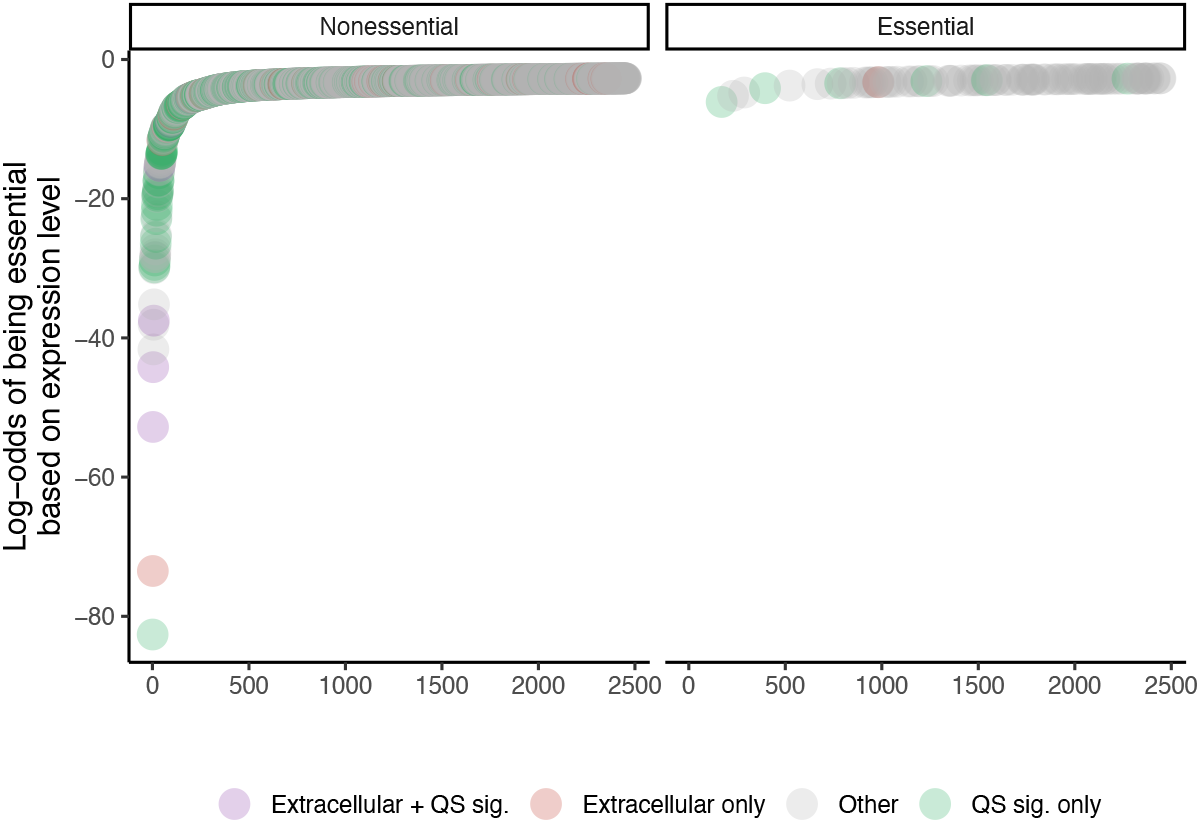
Extracellular and QS-regulated genes are enriched among genes with the lowest predicted probability of essentiality. Genes are ranked by their log-odds of being essential (predicted from absolute fold change using binomial logistic regression). Points are colored by group membership: genes that are both extracellular (PSORTb predicted localization) and significantly QS-regulated (microarray p<0.05, downregulated in the absence of QS signal; purple), extracellular only (red), significantly QS-regulated only (green), or neither (grey).

The ability of a phage gene to be complemented is likely a product of that gene’s role in the phage life cycle. For example, under different MOI conditions, genes that impact phage replication (manipulation of host factors to replicate phage genomes) will act differently than genes that impact phage exit (packaging and lysis) or phage adsorption. These difference could be viewed due to the impact that different scales of competition (local and global competition) will have on the gene.

To illustrate some key differences, we consider how phage genes can be affected across the phage life cycle; receptor-binding proteins cannot be complemented, in singly infected host cells within a generation, therefore mutants defective in adsorption should be depleted in Tn-seq data, in a single generation, regardless of the presence of other phages. De novo arising receptor mutants could of course be complemented by other phage present during lysis. By contrast, intracellular processes such as host manipulation, and lysis may be complemented under the same assumptions, allowing defective genomes to persist or even increase in frequency when coinfection is common or production of a public good is expensive or costly in some manner. To further differentiate between replication and exit genes we can consider burst size. If we assume a mutant of a replication (low MOI) will produce phage particles at a reduced rate, however, the subsequent burst size (the number of phages produced from a single infection) will be directly proportional to the phage production rate. Effectively, the number of phages produced inside the bacteria will be roughly equal to those that exit. A exit mutant, on the other hand, at a similarly low MOI, will have a different ratio; it is able to manipulate the host, increasing the number of phage particles inside but may lack any observed burst due to a defective lysis pathway.

To identify the role that gene sociality might play in masking gene essentiality we first examine the more well explored system of social genes in bacteria using data from a microarray under Quorum sensing induction and a bacterial Tn-seq experiment. We know which genes are typically considered social and even those we would expect to reduce in frequency from single mutant experiments. This analysis confirmed our assumption that in bacteria social genes are classified as less essential than expected. We then model phage mutant dynamics using stochastic simulations in which we simulate a Tn-seq like experiment where mutant genomes infect host cells across a range of MOIs and we track the abundance of progeny. Each gene has a fitness effect randomly chosen from a flat distribution. In this framework, assuming no mutation, the probability that a defective mutant is rescued depends on both the frequency of coinfection and the second randomly selected parameter, a social interaction (σ) that determines whether gene products are private (σ = 0) or fully public (σ = 1). Simulations reveal that private genes (σ = 0) reduce in frequency across MOI values based on their fitness effect, whereas socially complementable genes (σ > 0) can persist or increase in frequency as MOI rises. We extend this model to explore the consequences of phage lifecycle, where genes are separated to three life cycle compartments; Adsorption, Replication, and Exit. Using in-silico Tn-Seq experiments, we generate quantitative predictions for how gene frequency shifts should vary with MOI, how those shifts relate to phage lifecycle stages, and identify signatures of socially mediated complementation in phage populations.

## Methods Modelling

We developed a stochastic, agent-based simulation framework in R (version 4.3) to model transposon insertion sequencing (Tn-seq) experiments in phage. The simulation was designed to investigate how multiplicity of infection (MOI) interacts with gene-level complementation to distort apparent fitness measurements across distinct phage life cycle stages. All simulations were run using a saturated mutant library model in which every phage in the population carries a transposon insertion in exactly one gene, with insertion sites drawn independently and uniformly at random from the full gene set.

The simulated phage genome comprised 375 genes, each assigned two independent parameters: an intrinsic fitness effect and a complementation fraction. Genes were partitioned equally into three essentiality classes: 125 essential genes with a fitness effect drawn from a uniform distribution (Uniform[0.00, 0.10]), 125 moderate-effect genes (Uniform[0.10, 0.70]), and 125 near-neutral genes (Uniform[0.70, 1.00]). The fitness effect represents the relative phage burst size when the gene is disrupted, where a value of 0 indicates complete lethality and 1 indicates no fitness cost. We used numerous different distributions for the fitness including an approximated distribution from the Chan et al paper (SI X). Our results were highly robust regardless of fitness distribution or gene number (SI fig X). Therefore, for simplicity, we choose to focus on uniform distributions and 375 genes. Independently, genes were assigned to three complementation classes of equal size: low (Uniform[0.00, 0.10]), moderate (Uniform[0.10, 0.70]), and high (Uniform[0.70, 1.00]). The complementation fraction quantifies the proportion of the fitness defect that can be rescued when a coinfecting phage supplies a functional copy of the gene that is trans-acting. This could be consider a form of phenotype hiding [21]. Both essentiality class and complementation class were assigned independently, and genes within each class were assigned to life cycle stages by random shuffle. Our current framework does not capture the cost of producing a gene product, so our model does not capture cheating or cooperation, instead we assume a null position on the sociality of a gene, in reality behaviors that can be cheated on would increase in abundance beyond our predicted null model.

### Statistics

To investigate whether extracellular localization and quorum sensing (QS) regulation were predictive of gene essentiality in *P. aeruginosa* PAO1, we fitted a binomial logistic regression model with gene essentiality (as determined by transposon sequencing) as the binary outcome. In Turner et al essential genes were defined as any gene in which the expected proportion was significantly reduced below a log 2-fold reduction and a significance of 0.05 after correcting for multiple testing [6, 7]. We combined this with a microarray designed to detect QS controlled genes in PA [22]. Predictors included the absolute fold change in expression from microarray data (a continuous measure of QS regulation strength), predicted extracellular localization (binary), and their interaction term. Fold change and extracellular localization data were derived from a HSL condition microarray experiment, with analysis restricted to genes upregulated in the presence of QS signal (QS-induced genes). All statistical analyses were performed in R 4.3.2 using the base glm function with a logit link.

## Results

Most Tn-seq experimentation has been conducted in bacteria. The Gram-negative bacteria *Pseudomonas aeruginosa* has had both Tn-seq conducted (across a wide range of environments) and its social genes well characterized. Using these two data sources we show that genes that are controlled by quorum sensing (QS) regardless of localization are overrepresented as non-essential in Tn-seq data (**Fig 1**.). A logistic regression of absolute fold change from a microarray conducted on a Δ*lasIrhlI* strain of PAO1 supplemented with 15 µm of both 3-oxo-C12-HSL and C4-HSL was significantly negatively associated with the Tn-seq assessment of gene essentiality (Log odds estimate -1.42, p = 0.002). The more highly a gene was expressed under QS the more likely it was to be classified as non-essential. A similar analysis was performed on genes that were downregulated by QS. QS down regulated genes had a 2.1 folder greater probability of being classified as essential by TN-seq data (Log odds estimate = 0.75, p < 0.001).

Under typical lab conditions, (high MOI) coinfection of bacterial cells by phages is not exceptional but expected. For an infection, the probability that a bacterium receives at least two phages follows a Poisson distribution and can be calculated by 1 − e^−MOI^(1 + MOI). At an MOI of approximately 1.67, the threshold is crossed such that the majority of infected cells harbor more than one phage (**Fig 2**).

**Figure 2.**
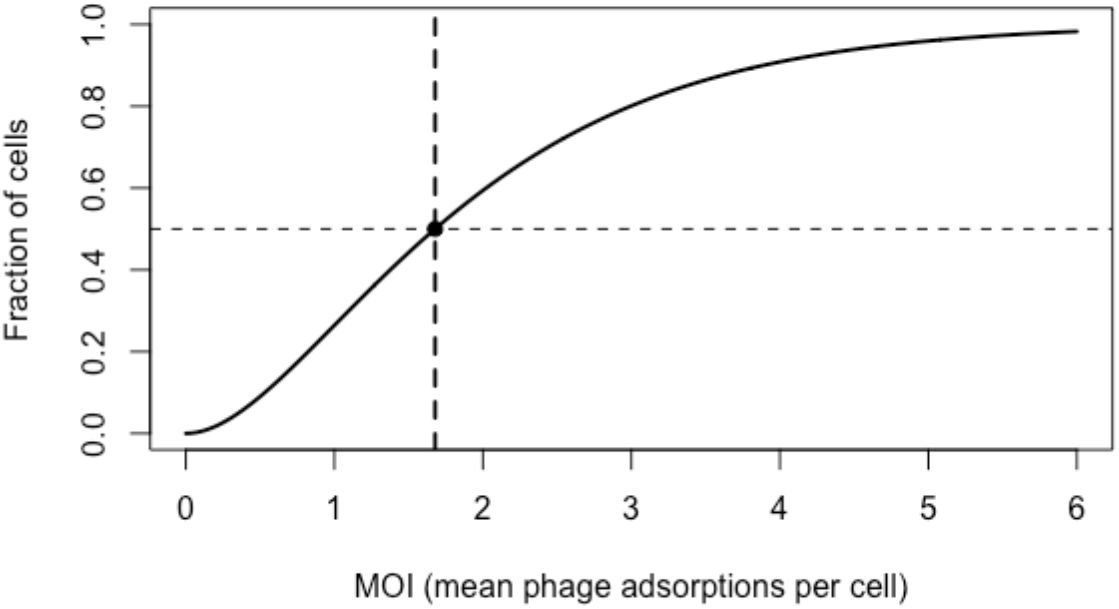
MOI tipping point of more than one phage per cell on average. As MOI increases the number of phage adsorptions increases. At an MOI of 1.67 more than 50% of the cells are infected with 2 phages. It is worth noting that not all cells are infected with 2 phages, just that the majority of cells have at least 2 phages.

Phage genes can be complemented in trans ([8-10]). Using a simple model assuming no mutation, only two classes of genes, and without fitness benefits gained by lacking a gene, we can demonstrate that under different MOIs, social genes (those that can be complemented) will increases in frequency as MOI increases (**Fig 3**). This model describes two or more phage genomes coinfecting the same host.

**Figure 3.**
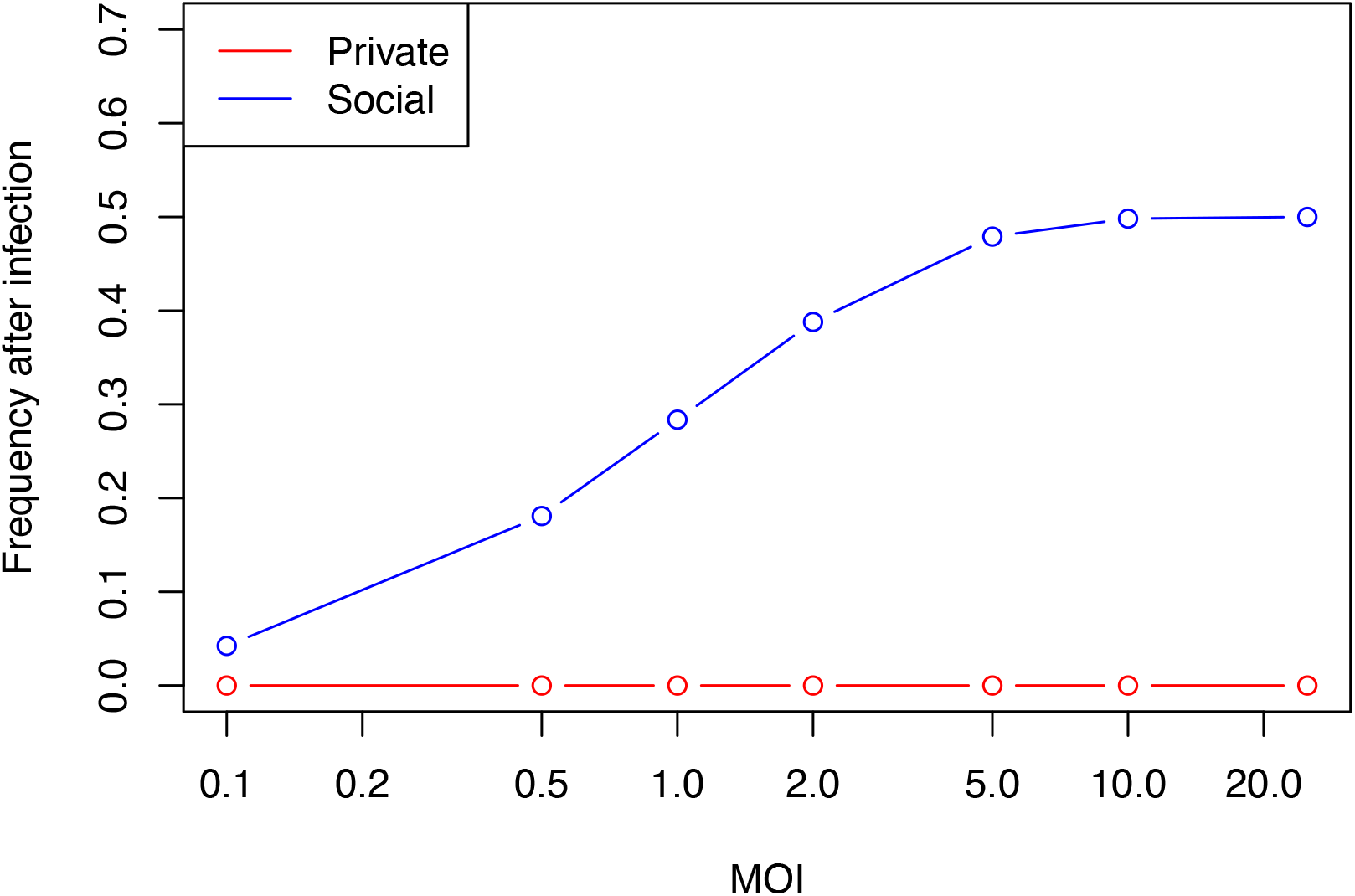
Abundance of private and social genes when coinfecting with a Wild Type phage. Social genes increase in frequency as the MOI increases. Under a simple model system, two genes were considered, a gene which could be complemented (social) and one that could not (private). As MOI increases the frequency of a social gene increases as it is being complemented by another genome.

Going beyond our simple model, we simulated a Tn-seq experiment in which 375 phage genes were randomly selected and assigned both a fitness score and a social factor (sigma), then allowed to infect hosts across a range of MOIs. Predictably, complementable genes, which had negative fitness effects at low MOIs, increased in abundance at higher MOIs (**Fig 4**). Under real world experimental conditions this would be seen as a reduction in the gene’s essentiality as MOI increases, effectively hiding the fitness impact of a gene.

**Figure 4.**
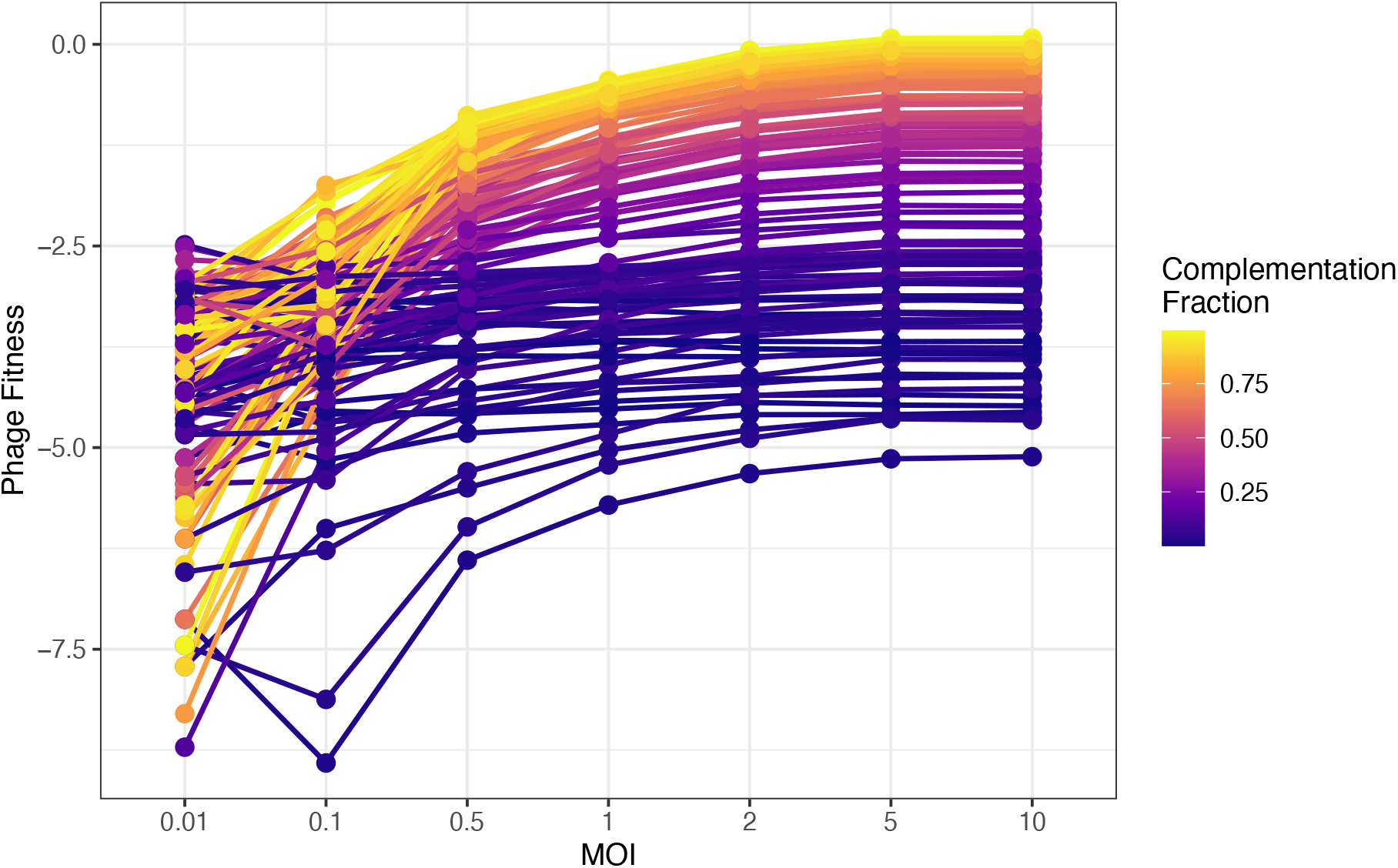
Using a Tn-Seq simulation, as MOI increases, the fitness (normalized abundance) increases if the gene can be complemented. Assigning 375 phage genes both a fitness effect and a complementation factor demonstrates that under increasing MOIs genes will increase in abundance when they can be complemented. Fitness is the difference in the (normalized log_2_) expected abundance of the gene.

Different parts of the phage life cycle may limit when genes can be complemented. We divided the phage genome into three key life cycle compartments; adsorption, which cannot be complemented, replication, which can be complemented, and exit, which can also be complemented, each received 125 genes. Genes required for adsorption likely cannot be complemented as adsorption happens before transcription occurs. While there may be instances where phages can interact at the adsorption step, we have simplified our assumptions here to exclude that possibility. As expected, our model shows that genes that can be complemented increase in abundance at higher MOIs (**Fig 5**).

**Figure 5.**
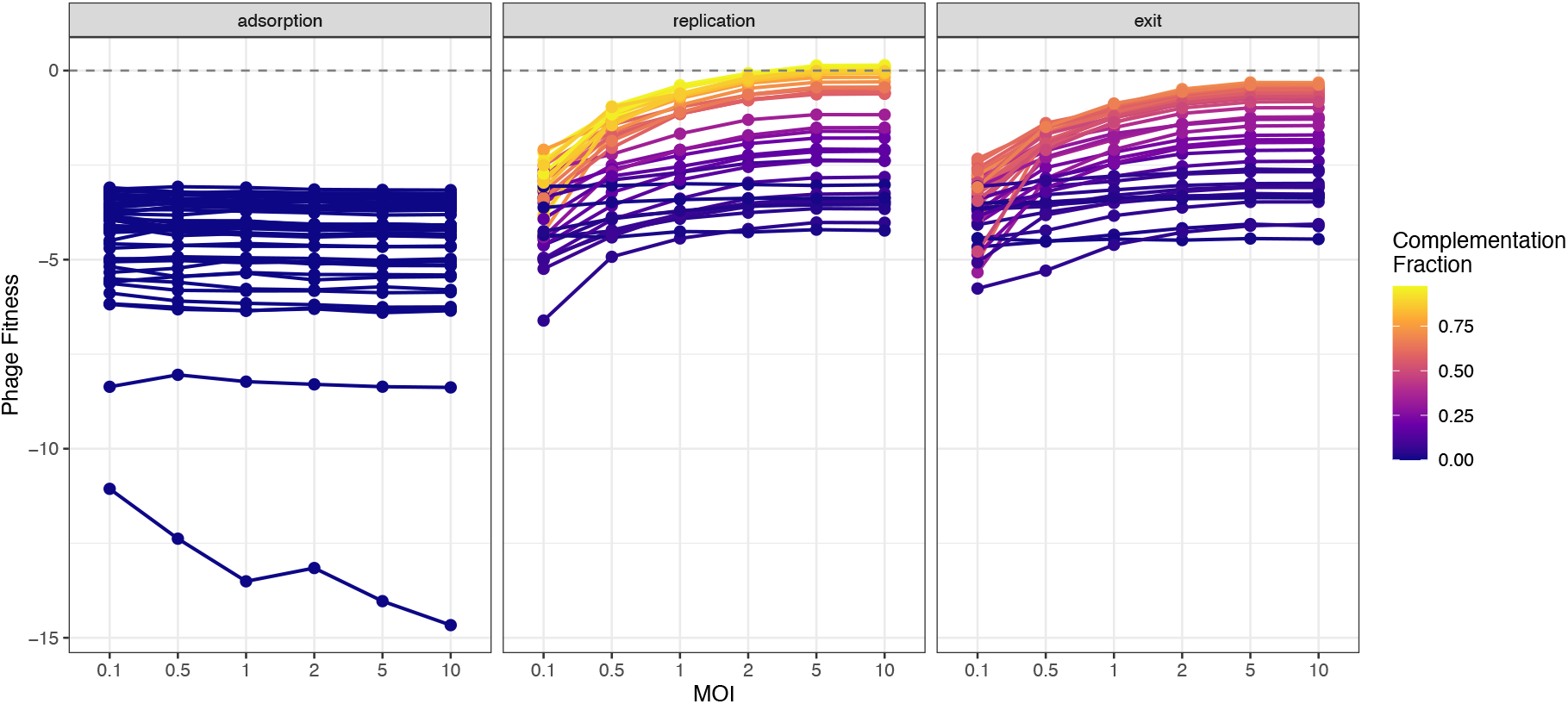
Essential genes across the three life cycle compartments – adsorption, host manipulation, and exit under a given MOI. Across a phages life cycle different genes are essential. Phage genes were separated into 3 different life cycle compartments with different abilities to be complemented and a distribution of fitness effects. Genes involved in adsorption could not be complemented and therefore did not increase in frequency in proportion to the MOI. Replication and exit genes could be complemented and therefore did increases in abundance under higher MOIs. Fitness is the difference in the normalized log_2_ expected abundance of the gene.

Although replication and exit genes behave similarly under a single coinfection round, a key distinction emerges when the scale of competition changes between rounds. To model this, we implemented a two-round infection design. In the first round, phages infect bacteria at a range of MOI values. The output phage pool from the first round with frequencies already shaped by complementation is then used as the input for a second round conducted at near-zero MOI (MOI = 0.01), where each phage effectively infects a cell alone. This second phase removes all complementation and exposes the single infection fitness of every mutant. This techniques has recently been used to explore mutations in phages by Meir et al [23]. Effectively we are testing the evolutionary life histories of the phages. The two rounds differ not only in complementation opportunity but also on a competitive scale. The first round involves local competition within cells, where social interactions (complementation and potential exploitation or cooperation) determine within-cell output. The second round involves global competition in which phage output is determined solely by intrinsic genotype, with no cheating or cooperation possible. An asymmetry between replication and exit mutants is revealed under these conditions. Replication mutants that were rescued in round one produce progeny at a reduced (but non-zero) intrinsic rate when alone in round two. Exit mutants, however, face a categorical failure: even if the phage genome replicates normally, without functional exit machinery, the cell cannot lyse, therefore no progeny is released. Exit mutant burst size is therefore zero when unaccompanied, regardless of the intrinsic fitness parameter value (**Fig 6**). There is of course the assumption that viral capsid is not carrying a protein that aids in lysis. This predicts that after a high-MOI first round, the output pool is enriched for exit mutants (rescued by complementation), but these mutants are then purged in the low-MOI second round. The Δ fitness score between rounds (the second-round score minus the first-round score) is negative for exit genes when the first round was conducted at high MOI. Adsorption genes regardless of MOI, are not subject to complementation and thus provide a control for our theory, assuming no phenotype mixing occurs.

**Figure 6.**
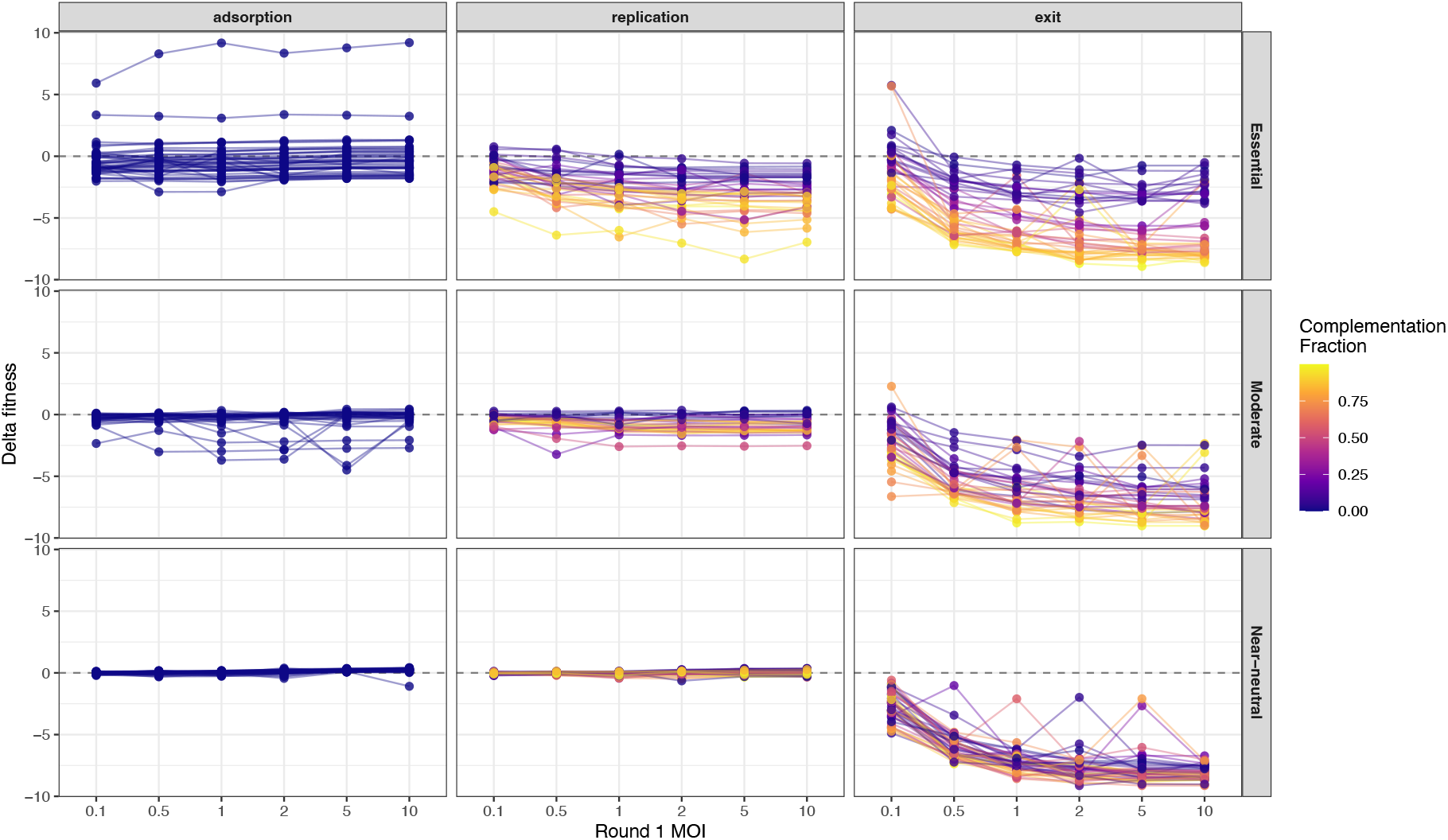
Two-round infection simulation across a range of first-round MOI values. Each phage from the first-round output pool infects a new host alone in the second round (MOI = 0.01). The delta score (round two minus round one) is shown for genes partitioned by stage and complementation fraction. Adsorption genes show delta ≈ 0 at all MOI values. Replication gene deltas are near zero to negative and scale with complementation fraction. Exit gene deltas are strongly negative owing to the zero-burst penalty imposed on unaccompanied exit mutants.

## Discussion

Sociality in nature is a broad topic; in recent decades socio-microbiology has increased in relevance with the acknowledgment of social interactions in single cell organisms ([16-18]). Phage genes have been shown to be social in empirical settings, with gene products shared among coinfecting genotypes in ways that mirror cooperation and exploitation in many viruses and other biological systems ([16, 18, 20]). Despite this, existing Tn-seq studies have not considered how social settings shape the apparent essentiality of genes, or genes required for phages to successfully complete their life cycles. Here we provide evidence for this being an understudied question by demonstrating that well described social genes in bacterial Tn-seq experiments are less essential than expected by random chance. There are two explicit hypotheses that could explain the significant reduction in QS controlled genes in the essential data. First, QS genes which are up regulated are never essential. Second, Tn-seq experiments underrepresent essential social genes due to complementation of the missing factor by the population. We can dismiss the first hypothesis as mutants of QS systems are known to have reduced fitness in the absence of Wildtype bacteria ([24]). QS mutants are able to cheat on bacteria producing products such as LasB protease ([25]). In the absence of Wildtype bacteria, these cheats cannot grow in a media which requires the production of LasB ([25]). However, as we have shown, in the Tn-seq data *lasB* mutants faced no fitness penalties, instead increasing in expected frequency ([6, 7]). Tn-seq studies of phage genes are likely to have similar issues, perhaps even amplified based on the MOI these experiments are conducted under. However, this may allow us to directly identify which genes are social in phages.

We extended our analysis to phage Tn-seq experimentation in silico. Our models show that the fitness scores produced by a Tn-seq experiment may not be the fixed properties of genes but can be affected by the social environment in which the experiment is conducted. In highly social contexts, those at high MOI, essential genes with high complementation fractions are systematically under-reported as essential as mutants are rescued by coinfecting genotypes, leading to the inflation of their frequencies in the output pool. While we did not manipulate genome size, the extent to which complementation inflates apparent fitness depends not only on MOI, but also on genome size. In a saturated mutant library, the probability that two randomly chosen phages carry disruptions in the same gene is 1/the number of genes. For a phage with a small genome of 20 genes, this probability is 5%, compared to just 0.2% for a 375-gene genome. This means that smaller phage genomes, paradoxically, experience less complementation-mediated rescue at a given MOI than larger genomes, because coinfecting phages are more likely to share the same disrupted gene and therefore cannot supply the missing product in trans. This can be recontextualized in evolutionary terms as their relatedness at any given loci is more likely to be 0 ([26]). We would predict that phages with small genomes might have fewer social genes after controlling for gene number and diversity. Our working model would be that smaller genomes are more likely to compete with less related individuals reducing the likelihood of cooperation evolving. However Polio and Influenza have been shown to have cooperative genes in the form of the capsid in Polio and both the capsid and replicase in Influenza and both have remarkable small genomes [27, 28].

To date experimental phage Tn-seq have deliberately used a very small MOI. However, by running multi-round experiments across MOIs it might be possible to expose stage-specific fitness effects. The two-round infection design we describe here provides a practical experimental strategy for distinguishing complementable from non-complementable genes, and perhaps for separating replication from exit defects. By conducting a first infection at high MOI to enrich complemented mutants, then passaging the output pool at near-zero MOI to strip away complementation, the resulting delta may partition genes along two independent axes: whether they can be complemented at all (reflecting life cycle stage and gene product biology), and how completely they are rescued when a coinfecting partner is present (reflecting the complementation fraction). We maintain that adsorption genes will produce delta scores near zero regardless of first-round MOI, serving as an internal negative control for this design. Exit genes, which produce the negative delta scores at high first-round MOI, are uniquely identifiable because their intrinsic burst size collapses to zero when unaccompanied. Experimentally, this could be tested by determining the abundance of phage inside bacteria before lysis and the production (burst size) post lysis. Naturally the difference may not be as clean cut as lysis or no lysis, ideally a population examination should reduce this error in measurement. Phage genes required for replication will result in a roughly equal abundances of phage particles internally to the size of the burst size. Again, using our modeling approach we can make predictions about the level of complementation expected in these categories.

The complementation dynamics we model are formally analogous to a number of social evolution dilemmas namely, public goods cooperation and local and global competition: a phage that produces a diffusible gene product and shares it with coinfecting genotypes is acting as a cooperator, while a mutant that benefits from that product without producing it is a cheater if the cooperator fitness is reduced ([29]). The fitness of cooperators and cheaters in this system is governed by the same inclusive fitness logic that applies to bacterial public goods games, with MOI setting the effective competition scales and potentially the relatedness of coinfectors ([18, 26]). At low MOI, selection operates within clonal groups, and cooperators are favored as the phage is only competing with its own genome and those directly replicated from it. At high MOI, mixing among genotypes increases non-cooperator frequency, consistent with the frequency-dependent dynamics predicted by kin selection theory ([30]). The key contribution of our model is to show that these social dynamics are a practical confound in an emerging genomic screen, and that they can be exploited, by deliberate MOI manipulation, to extract information about phage function that would otherwise be invisible to a single-condition Tn-seq experiment. This framework enables us to start mapping the sociality of phage genomes without having to know having to know much about these genomes or particular gene functions. The leveraging of Tn-seq tools in this way to better understand phage biology and their evolutionary dynamics are therefore a valuable and possibly unappreciated application of these tools.

Several limitations of the current simulation should be noted. First, we model complementation as a binary function of coinfector presence: a mutant is either alone (no rescue) or accompanied (partial or full rescue, according to the complementation fraction). In reality, the degree of rescue may scale with the number of coinfecting partners, with gene expression dynamics as we see in the bacterial QS and TN-seq data. There is excellent evidence that this might be the case with shared gene products in phages. Anti-CRISPR proteins have been shown to be more effective with increasing coinfections ([31, 32]). Second, our model does not include competition between phage genotypes within the same cell for limiting host resources, which could modify the effective burst size of complemented mutants in ways that depend on genotype composition. Rate vs. Yield mutation might become apparent in such circumstances [11, 18]. This would likely have a greater impact on genes that can be complemented as they would act as public goods, leading to no-cooperators increasing in frequency as they become cheats due to the negative fitness consequence on the cooperators ([18]). In this case a cheater genome would replicate to a greater percentage of all the phage from a single host.

While not a limitation, we choose to equally separate 375 genes among the compartments of phage life cycle genes, while phage genomes typically have asymmetric divisions in adsorption, replication and exit genes, this simplification does not impact our model to the best of our knowledge and instead allows us to study each compartment on equal terms. We also do not allow for mutation in our model, mutants are fixed. Finally, we used a uniform distribution to assign both complementation and fitness effects. We conducted the same simulation using a range of distributions including an approximation of those found in recent Phage Tn-seq data [3]. We found no difference in the outcome from changing this distribution and therefore used arguably the simplest distribution, uniform. Despite these simplifications, the core qualitative predictions, that complementable genes would be under-reported as essential at high MOI, that adsorption genes are immune to this effect, and that exit genes show a distinctive zero-burst phenotype when unaccompanied, are robust to the specific parameter values used and follow directly from the biology of the phage life cycle. Phage life cycles might also impact how easily phage genes are complemented. Phage Tn-seq data from Chan et al used the jumbo phage PhiKZ. This phage is known to make a phage nucleus. To the best of our knowledge, it is unknown if multiple phages can share a single nucleus or exclude each other and therefore might exclude products from other phage genes. The form of TN-seq experimentation we are describing might be able to determine that. Going further, we currently do not have a good understanding of which genes might act synergistically with each other or those that allow for cheating to occur. Phages mutants that cheat would exceed the expected frequencies from our models in terms of abundance but the experiments we are modeling would capture them.

Looking forward, coupling the simulation framework developed here with empirical Tn-seq data collected across an MOI gradient would provide a powerful approach to both empirically validate at a gene-by-gene level the sociality of phages and potentially allow for the annotation of phage gene function at scale. Genes whose fitness scores are stable across MOI would be classified as private (non-complementable), pointing to adsorption functions or cis-acting genomic elements. Genes whose scores rise with MOI would be flagged as social, with the slope of the MOI-fitness relationship providing an estimate of the complementation fraction. Genes that show the strongest delta scores in a two-round experiment would be prioritized as exit function candidates.

In summary, this approach of varying MOI in phage Tn-seq experiments could be applied to any lytic phage for which a transposon library can be constructed, providing a route to systematic functional classification and sociality classification of phage genes. Potentially more interesting is the idea that different phage life history and phage genome sizes could have a different selection pressure on the abundance of social genes.

## Acknowledgements

For funding, we thank Georgia State and the Cystic Fibrosis Foundation for the grant (GURNEY20F0-CI) and (GURNEY25I0) to J.G. We thank Asher Leeks, Iris Irby, Eli Mehlferber, Sam Brown, and Sean Czerwinski for their comments on the draft manuscript.

## Notes

### Competing Interest Statement

The authors have declared no competing interest.

